# Reviving ghost alleles: Genetically admixed coyotes along the American Gulf Coast are critical for saving the endangered red wolf

**DOI:** 10.1101/2021.12.18.473316

**Authors:** Bridgett M. vonHoldt, Joseph W. Hinton, Amy C. Shutt, Jennifer R. Adams, Lisette P. Waits, Kristin E. Brzeski

## Abstract

The last red wolves were captured along the Gulf Coast in 1980, where they hybridized with coyote, to establish the captive breeding population. However, red wolf ancestry persists in local coyotes and could be leveraged by genomic innovations to support species persistence. We assessed genomic ancestry and morphology of coyotes in southwestern Louisiana, and find they carried 38-62% red wolf ancestry acquired in the last 30 years, which is enriched on land with minimal coyote hunting. These coyotes were also similar in ancestry to canids captured in the 1970s that initiated the red wolf captive breeding program. Further, we reported that coyotes with higher red wolf ancestry are larger in size. Our findings evidence the importance of hybrids as a reservoir of endangered species ancestry for contemporary conservation efforts. Admixed genomes are at the forefront of innovative solutions, with red wolf survival a prime candidate for this new paradigm.

**Teaser:** Coyotes along the American Gulf Coast carry genetic variation critical for the survival of the endangered red wolf species.

## Introduction

The conservation of hybrids remains a contentious and pressing issue in conservation biology (*1*). As human activities such as anthropogenic mortality, habitat degradation, and translocations of organisms promote increased incidents of species hybridization and introgression (*2–4*), increased interest for a web-of-life framework has been considered when developing conservation strategies for imperiled species (*5*). For example, allowing for some limited level of gene flow between species may facilitate genetic rescue for small, inbred populations (*6*) by countering the negative consequences of small effective population sizes with the positive consequences of novel allelic combinations (*7,8*). Further, such genetic exchange can promote rapid evolutionary innovation and adaptation, particularly under a changing climate (*3,8,9*), which may be considered an untapped mechanism of conservation and preservation of genetic variation. However, the policy for the management of hybrids and admixed individuals is unclear with hybrids rarely offered legal protections, partly due to the difficulty of classifying and measuring the impact of hybrids on parental species and environments (*10*). Yet admixed genomes are a proven reservoir of novel genetic and phenotypic combinations upon which natural selection could act (*3*).

Genomic research can identify signatures of past genetic exchange (i.e., ghosts of introgression) in admixed genomes (*11*). Genetic traits once thought extinct can be rediscovered and potentially revived when innovative conservation practices are considered. While traditional practices remain critical for species persistence, new genomic technologies paired with extreme reproductive assistance, such as cloning and biobanking, can create new frontiers in conservation biology and hold new promises for species on the brink of extinction (*12–15*). Conservation practitioners are now supported with unprecedented technologies to construct clones or specific hybrid individuals that contain edited genomes that resurrect ghost variants and restore historic genetic variation. These pioneering methods create a new space where admixed individuals play an important role in species conservation as critical reservoirs of ghost genetic variation.

Here, we provide a timely study pertinent to the red wolf (*Canis rufus*), a critically endangered species endemic to the southeastern United States, and coyote (*C. latrans*), a species ubiquitous across North America. The survival of the red wolf could benefit from genomic technologies to bolster genetic variation as every extant red wolf is a descendant of 14 founders, which has severe demographic and genetic consequences (*16*). Red wolves and coyotes have hybridized both historically and contemporarily (*17–21*). Most notably, during the mid-20th century, the last known red wolf populations along the Mississippi River Basin were extirpated and the remaining wolves along coastal regions of eastern Texas and southwestern Louisiana began hybridizing with coyotes colonizing the region as wolf populations declined (*17,22,23*). Consequently, the United States Fish and Wildlife Service (USFWS) listed the red wolf as endangered and removed the last known individuals from the wild by 1980 to establish a captive breeding program as part of their Species Survival Plan (*24,25*).

Despite the disappearance of the red wolf, reports of wolf-like canids in rural regions of coastal southeastern Texas and southwestern Louisiana accumulated over the subsequent decades (*26–28*). Two recent independent studies substantiated these reports when red wolf ancestry was discovered in coyote populations occurring in eastern Texas and southwestern Louisiana (*27–29*). Further research has demonstrated this Gulf Coast region likely represents a focal region of red wolf ancestry that has persisted since red wolf extirpation in the 1970s (*21,29*). Although these studies lacked associated morphology, it is reasonable to hypothesize that higher red wolf ancestry resulted in the large-bodied *Canis* documented in Louisiana (*26*). A previous study identified introgression of putatively functional variation in the admixed genomes of canids of northeastern United States (*30*). Thus, given the variable phenotype and genomic ancestry observed across southeastern canids, we hypothesize a correlation with species-specific morphometrics, as measured by body size. These introgressed coyotes along the Gulf Coast states could represent a unique reservoir of previously lost red wolf ancestry, which has persisted in coyote genomes and could be critical for combating inbreeding in the genetically-limited extant captive red wolf population. The integration of morphology and genome ancestry would present a uniquely powerful tool for prioritizing the selection of individuals to boost long-term health of the critically endangered red wolf species.

Here, for the first time, we integrate genomic ancestry and morphology of coyotes living along the American Gulf Coast. We accomplished this by assessing red wolf ancestry in coyote populations along coastal southwestern Louisiana (hereafter “SWLA”) where red wolf and coyote hybridization occurred (*17,28,31*). By capturing coyotes in these admixed populations, we acquired both genomic and morphologic data to identify the quantitative thresholds by which one could prioritize animals for potential use in ongoing red wolf recovery efforts based on individual ancestry proportions combined with phenotypic traits such as body size. Although it is known that hybrids are intermediate in size to red wolves and coyotes (*17,26,32*), the correlation of body size and ancestry is not well documented. Therefore, we investigated the effects of autosomal and X-linked red wolf ancestry on coyote body size. We then consider landscape characteristics that likely supported high retention of red wolf ancestry in coyote populations without management, revealing land cover where red wolf ancestry is most resilient. We suggest hybrids are critical for defining what constitutes a red wolf, and admixed genomes will be pivotal in aiding red wolf conservation.

## Results

### Capture and collaring of Louisiana coyotes

We captured and radio-collared 26 coyotes (9 females, 17 males) from Cameron and Jefferson Davis Parishes of southwestern Louisiana between February 7th and May 6th of 2021. We collected a combination of blood and ear tissue from radio-collared coyotes following the approved IACUC protocol at Michigan Technological University (#1677987-2). We opportunistically sampled ear tissue from seven road-killed coyotes (1 female, 6 unknown) from SWLA and a single male SWLA coyote in a wildlife rehabilitation facility. Coyotes in SWLA had a general appearance intermediate that of western coyotes and red wolves of North Carolina (*26*) (Fig. 1).

**Fig. 1.**
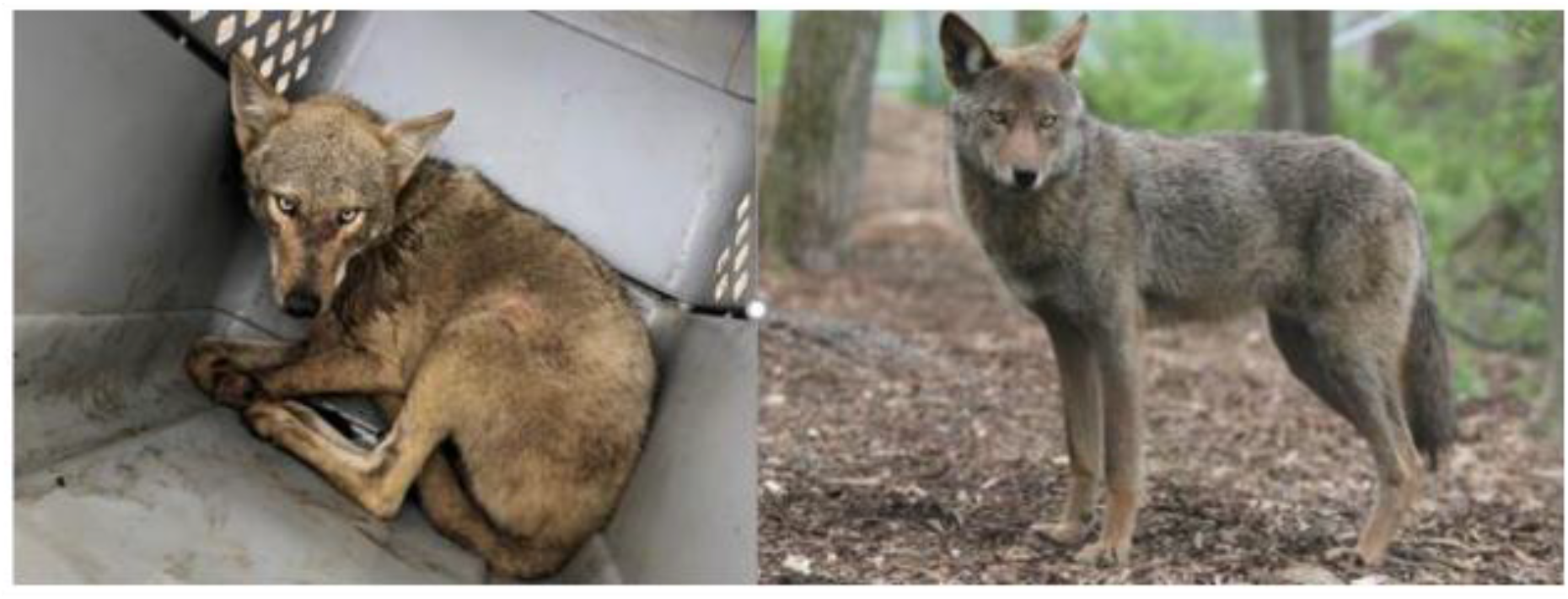
Comparison of coyote and red wolf. Coyote CL12928 (left panel) captured in southwestern Louisiana compared with a captive red wolf at the Wolf Conservation Center in New York (right panel, courtesy of Maggie Howell).

### Louisiana coyotes have genetic signals of reference coyotes and red wolves

Due to the high variability in the phenotype of SWLA coyotes (*26*), we obtained restriction-site associated DNA sequence (RADseq) data from 44 samples representing 34 unique coyotes from Louisiana and 10 red wolves from the North Carolina Non-essential Experimental Population (NCNEP, *33*) (Table S1). NCNEP red wolves were included for comparison given that they have experienced minimal introgressions from coyotes since reintroduction and could be genetically similar to canids along the Gulf Coast. We merged the genome-wide SNP genotype data with publicly available data from an additional 88 canids that represented several distinct reference lineages: 10 domestic dogs, 39 coyotes, 19 gray wolves, 10 eastern wolves, and 10 captive red wolves from the Species Survival Plan (SSP) population (*21,27,34*) (Table S1). After extensive data filtering, we retained 130 canids and 59,788 SNP loci out of a total of 199,888 catalogued variants. Additional further filtering for linkage and Hardy-Weinberg Equilibrium (HWE) deviations established a subset of 41,309 SNP loci that we designated as statistically neutral and unlinked. A principal component analysis (PCA) revealed the expected clustering of each canid reference lineage, while the NCNEP red wolves clustered tightly with the SSP reference red wolves and the Louisiana coyotes spanned two PC2 clusters of red wolves and coyotes (Fig. 2). Given the lack of variation among the NCNEP red wolves, they were included with the SSP reference red wolves for downstream analyses.

**Fig. 2.**
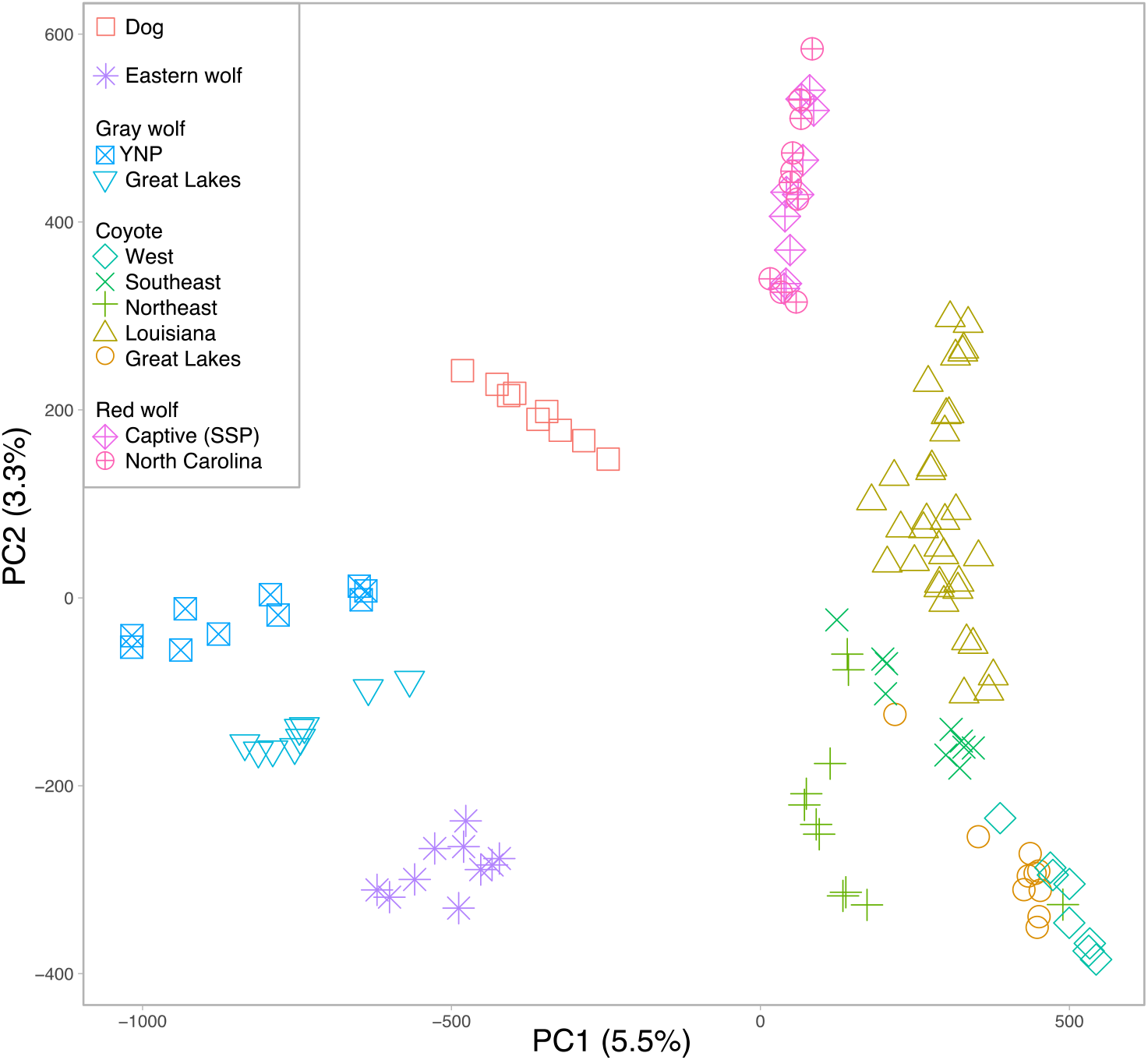
A principal component analysis of 130 canids genotyped at 41,309 SNP loci. The percent of variation explained for each axis is provided in the parentheses. (Abbreviations: SSP, Species Survival Plan; YNP, Yellowstone National Park)

### Coyotes of southwestern Louisiana carry high red wolf ancestry with recent admixture dates

To investigate the degree and geographic extent to which these coyotes may be a reservoir for lost red wolf genetic variation, we inferred red wolf ancestry proportions for 31 Louisiana coyotes across 59,788 SNP loci. We found that these individuals displayed variable red wolf ancestry proportions across the autosomes (mean±s.d.=0.38±0.2) and X chromosome (0.62±0.3) (Table 1). Given our limited geographic access for sample collection, we have an enrichment of genetic representation within Cameron Parish; however, concordant with past findings, this parish also contained the highest red wolf ancestry proportions (min-max: autosomes=0.18-0.69, X chromosome=0.18-1.0) with the most recent estimated admixture timing (autosomes=20 years, X=24 years) (*21,29*) (Fig. 3A, Table 1). The other three Louisiana parishes collectively analyzed (Jefferson Davis, Iberville, and East Baton Rouge) were significantly lower in average red wolf ancestry (autosomes=0.21, *t*-test *p* 2×10^-5^; X=0.42, *p*=0.0035) with older admixture timing (autosomes=25 years, *p*=0.002; X=26 years, *p*=0.2554). We visualized the location of ancestry blocks across the chromosomes of a coyote with the lowest red wolf (RW) proportions (sample CL12938), alongside the coyote with the highest RW proportions (sample CL12939). We found that such fragments are frequently in the heterozygous state for the low RW content coyote, while the high RW content coyote’s genome carries a substantially higher frequency of homozygous red wolf blocks (Fig. 3B).

**Fig. 3.**
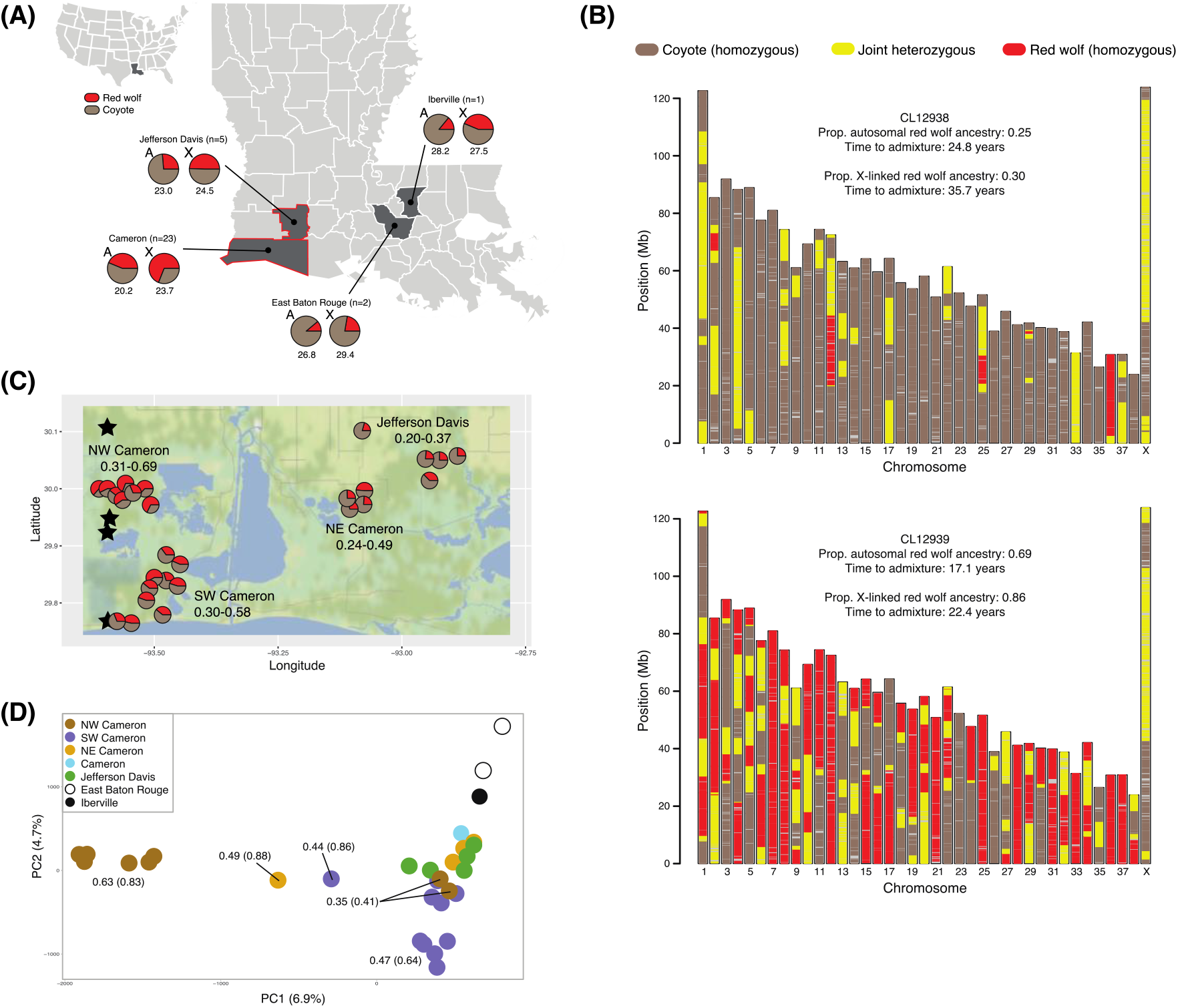
Genomic ancestry of 31 Louisiana coyotes. **(A)** Ancestry was inferred across 59,788 SNP loci across autosomes (A) and the X chromosome (X) with respect to two reference lineages: 39 reference coyotes and 10 reference red wolves from the captive Species Survival Plan population. The average timing of admixture is provided (in years) below the pie charts. The parishes outlined in red are enlarged in panel **(B)** Chromosomal plots of ancestry fragments for two Louisiana coyotes. Fragments across each of the 38 autosomes and X chromosome are color coded with respect to ancestry state. These two individuals were selected to display the lowest/oldest and highest/most recent red wolf proportions (CL12938 and CL12939, respectively). Sample identity, ancestry proportions, and admixture timing estimates are provided above each plot. **(C)** with higher resolution spatial details of autosomal proportions of red wolf ancestry (min-max) for each individual coyote with latitude/longitude data (see Table S1). Stars indicate the geographic locations of where four red wolf SSP founders originated. Sample sizes (n) and parish names are provided. **(D)** The PCA of 31 Louisiana coyotes revealed PC1 as an axis of red wolf ancestry (average autosomal red wolf proportions are provided for each color symbol with X chromosome proportions in parenthesis in the PCA). The percent of variance explained by each component is provided on each axis. (Abbreviations: NE, northeast; NW, northwest; PC, principal component; Prop., proportion; SW, southwest)

**Table 1.** Proportion and timing (in years) of red wolf genomic ancestry for 31 coyotes captured and sampled in southwestern Louisiana. Genomic ancestry was inferred across 59,788 SNPs genotyped in 31 coyotes from Louisiana with respect to 39 reference coyotes and 10 reference red wolves from the captive Species Survival Plan population. (Abbreviations: prop., proportion)

| Sample | Louisiana Parish | Prop. of red wolf ancestry: |  | Timing of red wolf admixture: |  |
| --- | --- | --- | --- | --- | --- |
|  |  | Autosomal | X-linked | Autosomal | X-linked |
| CL12923 | Cameron | 0.629 | 0.561 | 16.5 | 35.7 |
| CL12924 | Cameron | 0.678 | 0.957 | 14.7 | 0.0 |
| CL12926 | Cameron | 0.635 | 0.883 | 15.1 | 26.2 |
| CL12927 | Cameron | 0.573 | 1.000 | 17.5 | 0.0 |
| CL12928 | Cameron | 0.394 | 0.632 | 20.8 | 34.9 |
| CL12929 | Cameron | 0.595 | 0.723 | 16.9 | 20.2 |
| CL12930 | Cameron | 0.487 | 0.877 | 19.7 | 31.3 |
| CL12931 | Cameron | 0.241 | 0.901 | 25.4 | 37.9 |
| CL12932 | Cameron | 0.286 | 0.695 | 23.1 | 25.5 |
| CL12933 | Cameron | 0.254 | 0.526 | 24.9 | 16.3 |
| CL12935 | Jefferson Davis | 0.374 | 0.693 | 19.5 | 28.6 |
| CL12936 | Jefferson Davis | 0.247 | 0.314 | 20.7 | 18.8 |
| CL12937 | Jefferson Davis | 0.238 | 0.475 | 24.0 | 10.8 |
| CL12938 | Jefferson Davis | 0.249 | 0.296 | 24.8 | 35.7 |
| CL12939 | Cameron | 0.693 | 0.860 | 17.1 | 22.4 |
| CL12940 | Cameron | 0.312 | 0.184 | 21.6 | 18.0 |
| CL12973 | Cameron | 0.467 | 0.889 | 18.4 | 20.5 |
| CL12974 | Cameron | 0.300 | 0.911 | 23.6 | 25.0 |
| CL12975 | Cameron | 0.351 | 0.525 | 21.0 | 24.4 |
| CL12976 | Cameron | 0.435 | 0.860 | 19.8 | 16.8 |
| CL12977 | Cameron | 0.579 | 0.311 | 19.3 | 18.9 |
| CL12978 | Cameron | 0.391 | 0.868 | 21.0 | 35.4 |
| CL12979 | Cameron | 0.451 | 0.501 | 19.8 | 27.0 |
| CL12980 | Cameron | 0.389 | 0.743 | 20.9 | 31.8 |
| CL12981 | Jefferson Davis | 0.207 | 0.691 | 26.1 | 28.5 |
| CL12982 | Cameron | 0.446 | 0.777 | 19.1 | 19.5 |
| CL12983 | Cameron | 0.312 | 0.447 | 22.6 | 38.3 |
| CL13003 | East Baton Rouge | 0.098 | 0.336 | 26.5 | 24.2 |
| CL13004 | East Baton Rouge | 0.124 | 0.110 | 27.2 | 34.5 |
| CL13005 | Iberville | 0.141 | 0.437 | 28.2 | 27.5 |
| CL13006 | Calcasieu | 0.182 | 0.249 | 26.6 | 18.1 |

### Regional discovery of land preserves that lack hunting have highest red wolf ancestry

Of particular interest are the 28 coyotes sampled from Cameron and Jefferson Davis Parishes (Fig. 3A). Land cover across the region changes considerably with increasing distance from the Gulf Coast shoreline which was composed of a complex mosaic of saline to intermediate marsh zones in southeastern Louisiana (*35*). Much of the landscape in which we sampled coyotes had limited hunting access. For example, all 26 coyotes that we captured and radio-collared were of healthy weight and all mortality rates were both relatively low (N=4) and accounted for (vehicle collisions, N=2; capture myopathy, N=1; unknown cause, N=1). Coyotes with the highest red wolf ancestry were sampled in northwestern Cameron Parish (autosomes=0.56, X=0.73) on a private ranch that prohibited hunting and trapping of wildlife, followed by southwestern Cameron Parish (autosomes=0.41, X=0.68) on Sabine National Wildlife Refuge and corporate oil holdings with limited public access, northeast Cameron Parish (autosomes=0.32, X=0.75) on Cameron Prairie National Wildlife Refuge and surrounding private lands that permitted hunting, and in Jefferson Davis Parish (autosomes=0.26, X=0.49) on private land with active coyote control around its exotic hunting preserve (Fig. 3C). Chromosomal fragments of red wolf ancestry in the coyotes of northwestern Cameron Parish were the most recently acquired (autosomes=17.5 years, X=19.7 years), and again follows the same trend with older admixture time estimates with decreasing ancestry proportions (southwestern Cameron Parish: autosomes=20.5, X=25.8; northeast Cameron Parish: autosomes=23.3, X=27.7; Jefferson Davis Parish: autosomes=23.0, X=24.5).

We conducted a PCA of the 31 Louisiana coyotes and found that PC1 was negatively correlated with the average autosomal red wolf ancestry for each geographic origins of the samples (r=-0.844), and to a lesser degree ancestry on the X chromosome (r=-0.517) (Fig. 3D). We also find the continued support that coyote populations represent a mosaic of individuals with tremendous inter-individual variation in red wolf ancestry proportions, exemplified by the coyotes from northwestern Cameron Parish. Although this geographic cluster of samples contains individuals with the highest estimated red wolf ancestry, there are two with lower estimates and clusters with similar ancestry proportions on the PCA (Fig. 3D). Such variation highlights the need for rapid genome-level scans of populations critical for conservation practitioners and recovery programs.

The average longest homozygous red wolf ancestry blocks were carried by coyotes in northwestern Cameron Parish (56.2±49.4 Mb), relative to the other geographic clusters within the Parish (northeast=18.7±16.9 Mb, southwest=3.0±2.2Mb) and in the neighboring Jefferson Davis Parish (12.1±6.7Mb) (Table 2). This trend, however, is predominantly driven by three outlier individuals with extremely long homozygous red wolf ancestry blocks (87.2-122.6 Mb). The ratio of homozygous red wolf to coyote ancestry block sizes also revealed that coyotes in northwestern Cameron Parish had red wolf ancestry blocks 3.5 times longer than their homozygous coyote block sizes (56.2 Mb and 16.2 Mb, respectively). Coyotes in northeastern Cameron Parish carried the next longest red wolf ancestry blocks, 1.6 times longer than homozygous coyote blocks (18.7 Mb and 11.7 Mb), in addition to this region exhibiting older admixture time estimates relative to the northwestern Cameron region.

**Table 2.** Autosomal ancestry block sizes (in Mb) for 31 coyotes captured and sampled in southwestern Louisiana. Average block sizes for each ancestry state per coyote. Joint is defined as the heterozygous ancestry state. (Abbreviations: NE, northeast); NW, northwest; SW, southwest)

| Sample | Louisiana Parish | Coyote | Joint | Red wolf |
| --- | --- | --- | --- | --- |
| CL12923 | Cameron (NW) | 16.9 | 24.2 | 122.6 |
| CL12924 | Cameron (NW) | 9.0 | 25.7 | 122.6 |
| CL12926 | Cameron (NW) | 8.7 | 3.9 | 47.6 |
| CL12927 | Cameron (NW) | 14.9 | 7.8 | 16.0 |
| CL12928 | Cameron (NW) | 36.7 | 3.1 | 4.0 |
| CL12929 | Cameron (NW) | 9.1 | 26.6 | 87.2 |
| CL12930 | Cameron (NE) | 18.6 | 7.2 | 5.0 |
| CL12931 | Cameron (NE) | 23.5 | 0.7 | 3.1 |
| CL12932 | Cameron (NE) | 4.2 | 9.8 | 34.2 |
| CL12933 | Cameron (NE) | 0.5 | 0.3 | 32.4 |
| CL12935 | Jefferson Davis | 13.6 | 0.7 | 11.4 |
| CL12936 | Jefferson Davis | 6.7 | 8.2 | 23.7 |
| CL12937 | Jefferson Davis | 2.3 | 1.5 | 9.1 |
| CL12938 | Jefferson Davis | 30.8 | 46.3 | 8.3 |
| CL12939 | Cameron (NW) | 9.3 | 2.8 | 46.3 |
| CL12940 | Cameron (NW) | 24.9 | 0.4 | 3.6 |
| CL12973 | Cameron (SW) | 13.3 | 20.1 | 5.9 |
| CL12974 | Cameron (SW) | 20.9 | 1.2 | 2.5 |
| CL12975 | Cameron (SW) | 28.8 | 30.8 | 0.7 |
| CL12976 | Cameron (SW) | 5.1 | 1.1 | 2.8 |
| CL12977 | Cameron (SW) | 32.1 | 4.4 | 0.6 |
| CL12978 | Cameron (SW) | 10.7 | 8.0 | 5.1 |
| CL12979 | Cameron (SW) | 2.2 | 31.8 | 1.6 |
| CL12980 | Cameron (SW) | 33.6 | 0.6 | 3.1 |
| CL12981 | Jefferson Davis | 7.6 | 16.2 | 7.8 |
| CL12982 | Cameron (SW) | 10.1 | 18.8 | 6.9 |
| CL12983 | Cameron (SW) | 31.4 | 0.2 | 1.0 |
| CL13003 | East Baton Rouge | 13.9 | 11.4 | 6.0 |
| CL13004 | East Baton Rouge | 13.9 | 1.6 | 9.4 |
| CL13005 | Iberville | 15.4 | 13.0 | 10.4 |
| CL13006 | Calcasieu | 26.4 | 1.9 | 2.9 |

### Morphology and red wolf ancestry

We correlated body size of coyotes with red wolf ancestry estimates and found that coyotes with higher red wolf ancestry were on average heavier animals (Fig. 4). Coyote body mass was positively correlated with autosomal red wolf ancestry (β=5.95, SE= 2.77, *p*=0.044, R^2^= 0.25), and while the trend was similar with X-linked red wolf ancestry, it was not significant (β=1.54, SE= 1.99, *p*=0.45, R^2^= 0.11).

**Fig. 4.**
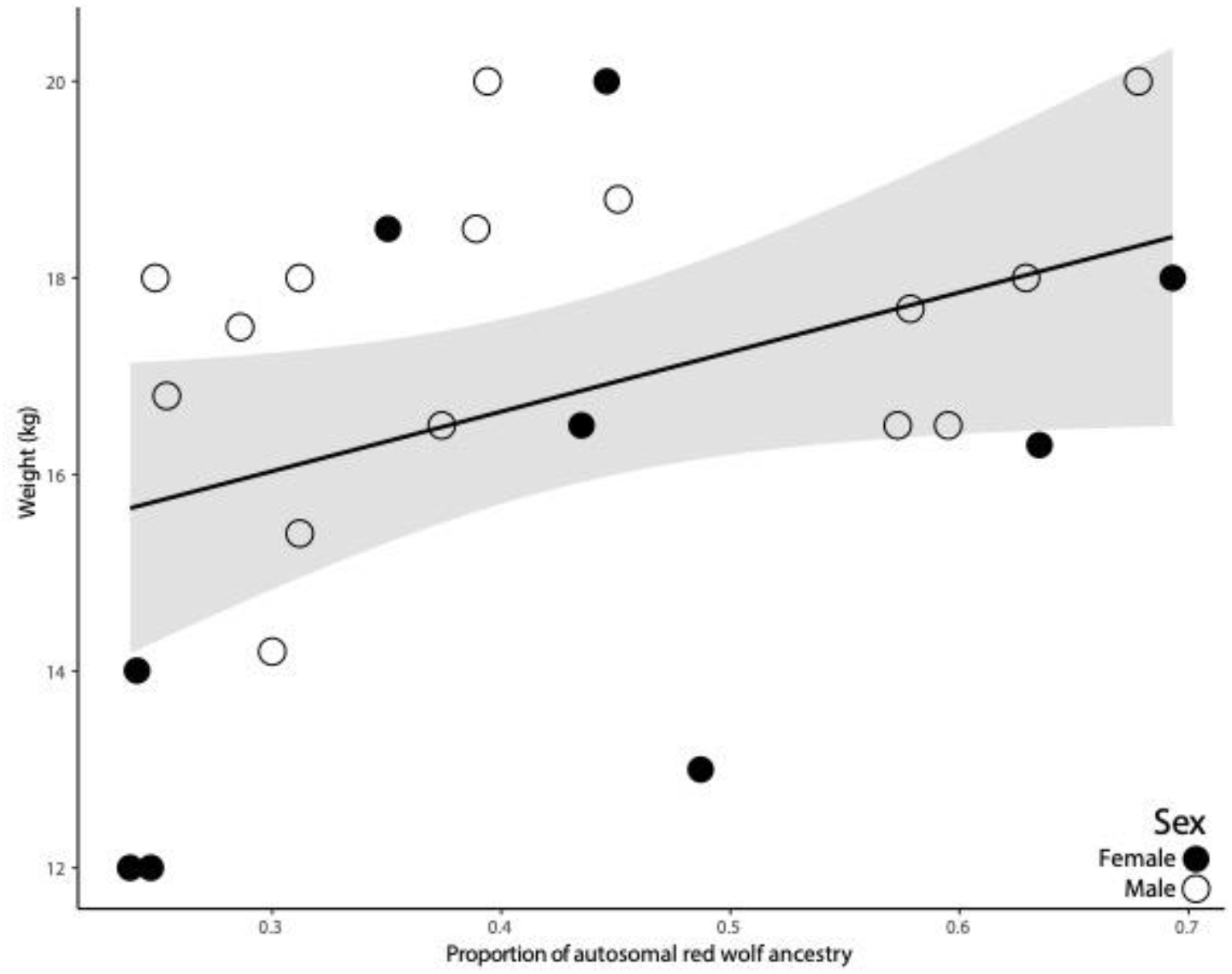
Body size and autosomal red wolf ancestry. Association between autosomal red wolf ancestry proportions and body size, as measured as weight (kg), for 24 live-captured Louisiana coyotes with ancestry estimates. The solid line is the fitted regression line and the grey shaded area represents SE of the linear model (β=5.95, SE= 2.77, *p*=0.044, R^2^= 0.25).

### Signatures of red wolf genetic variation in SWLA coyotes are similar to Texas canids from the 1970s

We included genotype data from 10 Texas canids sampled during the early 1970s capture efforts to identify the red wolf founders (Table 3). We annotated and genotyped 47 reference canids (coyote=37, red wolf=10), nine canids from 1970s, and 31 coyotes from SWLA for 45,994 SNPs after filtering for minor allele frequency, missingness, genotype correlation, and deviations from Hardy-Weinberg equilibrium. We found that several SWLA coyotes cluster in PC space proximal to canids from the 1970s capture events (Fig. 5A), with a maximum likelihood model-based approach discovering that coyotes from Cameron Parish have similar proportion membership values to red wolf cluster as the 1970s canids at *K*=4 (6% and 9.2%, respectively), relative to the other samples SWLA Parishes (<1%) (Fig. 5B). Such a trend holds for other partitions (*K*=3: 23.6%, 64.1%, and 32.6%; *K*=5: 8.3%, 5.8%, and <1%).

**Fig. 5.**
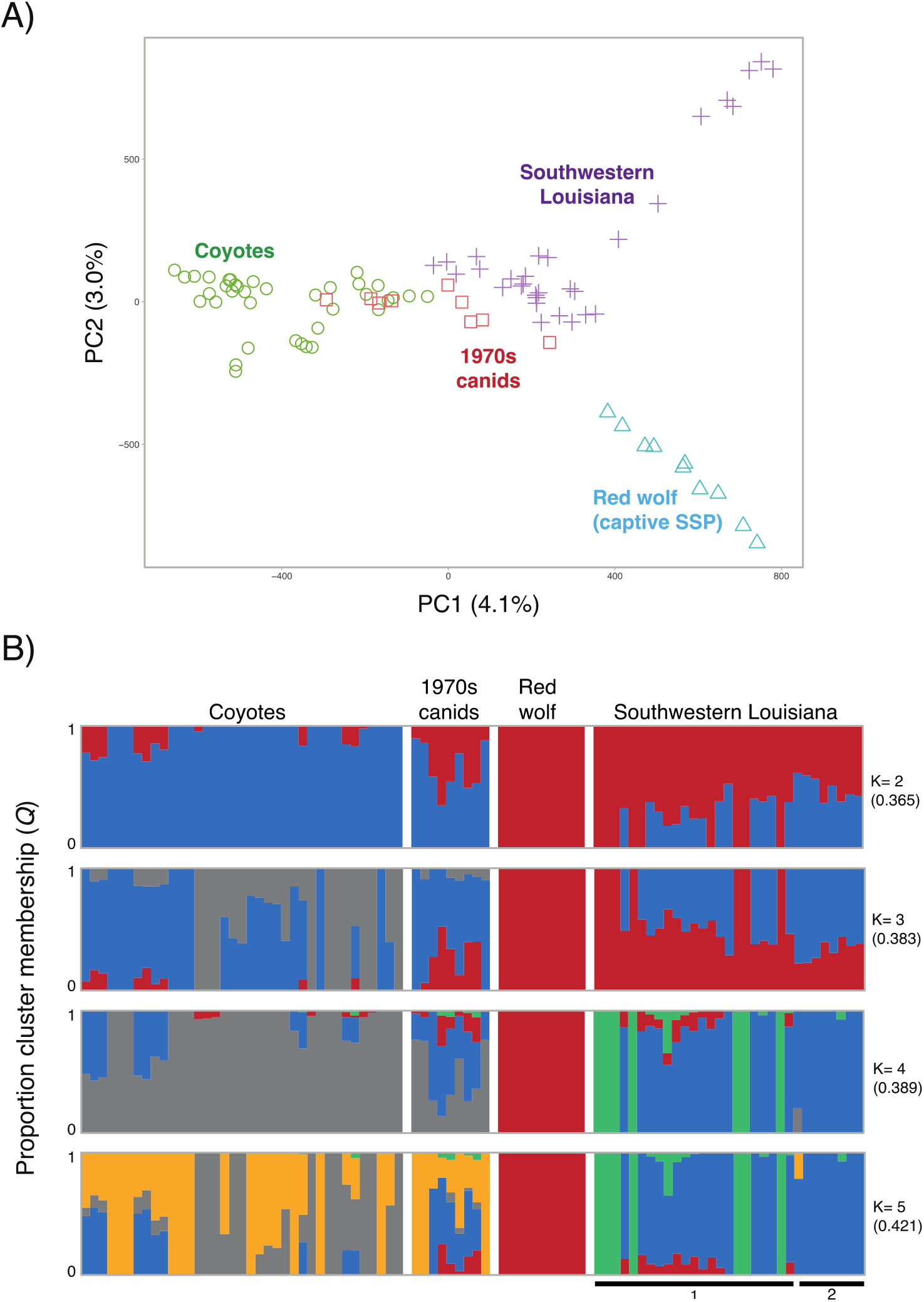
Genetic variation patterns of southwestern Louisiana coyotes with respect to Texas canids captured during the 1970s across 45,994 unlinked and neutral SNPs. **A)** A principal component analysis of 87 canids, and **B)** maximum likelihood cluster analysis for the best fit partition (*K*=2) and additional subsequent partitions (*K*=3-5). The cross-validation (cv) error per partition is given in the parentheses. The solid bar below the graph indicates the Parish of sample origins in Louisiana (1: Cameron Parish; 2: East Baton Rouge, Iberville, and Jefferson Davis Parishes).

**Table 3.** Sample information for 10 Texas canids from the 1970s red wolf capture efforts. Red wolf ancestry proportions are from previously published genome analyses (*21*).

| Sample ID<br>(County in Texas) | Red wolf ancestry<br>proportions | Collection<br>Date |
| --- | --- | --- |
| 70-TX-01 (Webb) | 0.02 | 3/22/1976 |
| 70-TX-02 (Webb) | 0.02 | 3/23/1976 |
| 70-TX-03 (Jefferson)* | 0.11 | 1/24/1976 |
| 70-TX-04 (Jefferson)* | 0.52 | 1/25/1976 |
| 70-TX-05 (Harris)* | 0.19 | 1/4/1976 |
| 70-TX-06 (Montague) | 0.02 | 11/21/1975 |
| 70-TX-07 (Brazoria)* | 0.03 | 2/2/1975 |
| 70-TX-08 (Liberty)* | 0.27 | 7/31/1975 |
| 70-TX-09 (Brazoria)* | 0.29 | 5/7/1975 |
| 70-TX-10 (Webb) | 0.02 | 3/23/1976 |

\*Counties geographically adjacent to or are actual red wolf SSP founder source locations.

## Discussion

Coastal SWLA is a particularly important locality for assessing red wolf-coyote hybridization as it was: 1) the last known area occupied by red wolves prior to their extirpation from the wild in 1980 (*17,31*); and 2) one of the first regions in the eastern United States to be colonized by coyotes (*36*). Indeed, we observed a range of red wolf autosomal ancestry (10-69%) in coyotes along coastal SWLA estimated to have occurred in the past 30 years. Coyotes with the longest and oldest contiguous chromosomal fragments of red wolf ancestry were found in the remote and isolated wetlands of Cameron Parish. In particular, coyotes with the greatest red wolf ancestry were on the FR Ranch of the Moore-Odom Wildlife Foundation, a property that has prohibited hunting and trapping of wildlife since 1911 (B. Bruce, *personal communication*). Our findings suggest that areas with reduced lethal management allow for the persistence of red wolf ancestry in coyote populations. Given that anthropogenic mortality of red wolves promotes wolf-coyote hybridization (*37–39*), we were not surprised that red wolf ancestry was greatest in coyotes residing in such isolated areas that afforded reduced exposure to lethal control.

We found a positive correlation between coyote body size and autosomal red wolf ancestry, and observed no coyotes with >50% autosomal red wolf ancestry that weighed less than 16.3 kg. The X chromosome has a more complex mode and history upon which natural selection can act. We find an enrichment of higher red wolf ancestry on this sex chromosome and suspect given more data, there is possibly sex-based differences in demography, life history, and fitness. Albeit a small sample size, our findings indicate that the contribution of large-bodied coyotes to the SWLA population is acquired through the inheritance of autosomal red wolf ancestry. Phenotypic characters such as body size and pelage color are helpful to identify hybrids and introgressed coyotes (*32,36*). For example, previous research has reported that only red wolf pups that had not achieved adult-like body sizes, rather than juvenile and adult red wolves, were confused for hybrids (*32*). They noted that hybrids were more similar to coyotes and no specific morphological character, except intermediate measurements and appearance, were used to differentiate hybrids from coyotes. Similarly, coyotes in SWLA that were highly introgressed with red wolf ancestry were phenotypically often more coyote-like than red wolf-like suggesting that genomic analyses using various genes are necessary to evaluate the extent of wolf introgression in the region’s coyote population.

Population surveys that employ genomic analyses will be critical to characterize the dynamics of the red wolf-coyote hybrid zone in the southeastern United States. Currently, it appears that remnants of the hybrid zone are now confined to isolated wetlands along the Gulf Coast that are considerably isolated from adjacent southeastern coyote populations. Our findings indicate that admixture occurred as recently as 20 years ago in an area that the USFWS Red Wolf Recovery Program re-surveyed for red wolves during the early 1990s (USFWS *unpublished data*). Although red wolves were not rediscovered by USFWS biologists, due to logistical difficulties when trapping in remote, isolated areas, biologists reported that they observed large canids and suggested that red wolves may have still existed in the region at the time (USFWS *unpublished data*). If our estimates of admixture are correct, our findings provide evidence that USFWS biologists were correct in their assessment that some red wolves may have persisted into the early 1990s.

### Importance of the admixture zone

Coastal SWLA represents a complex admixture zone and reservoir of presumed lost red wolf genomic variation as it persists in admixed coyote genomes. Further, given the enrichment of red wolf variation currently documented in this region, we suggest SWLA should be prioritized as a potential site for a future red wolf reintroduction. This natural occurrence of endangered genetic variation provides a redundant conservation design, supporting the SSP red wolf breeding efforts while providing a redundant and independent effective population. Our study presents one of the first connections between the red wolf phenotype and genomic estimates of red wolf ancestry. Building upon previous work that clearly supports red wolf-specific traits (*32,35,39*), we have documented a positive correlation between such traits and genomic ancestry in coyotes of the historic red wolf admixture zone (*17*).

Conservation practitioners are eager to implement innovative conservation strategies that incorporate functional genomic variation. Although we report the positive correlation between phenotype and ancestry proportions, it remains unclear which genomic regions are crucial for maintaining the red wolf phenotype. This challenge is driven by the historic distribution of red wolves across a diverse range of habitats, compounded by the rapid rate at which their historical landscape was permanently altered by European colonization and anthropogenic activity. We expect that, with further modeling of quantitative morphometrics and genomic variation, adaptive ancestry blocks can be linked to specific traits that are central to red wolf fitness, behavior, and ecology.

A final challenge presented in the admixture zone is species assignment. When a coyote is estimated to carry a predominant estimate of red wolf genomic ancestry proportion (i.e. >50%), we argue that such individuals are a crucial component for the persistence of the endangered red wolf. We have the tools to integrate ancestry estimates with ancestry block metrics (e.g. length, identity) to estimate the timing of which such events occurred. We encourage conservation practitioners to go beyond species concepts and pioneer a vision that leverages admixture to provide endangered genomes with the best possible probability for survival in our rapidly changing world.

### Conservation strategies

Species recovery plans have traditionally been organized around the model-based population viability analysis (PVA) to outline measurable recovery criteria (*40*). Challenges to such PVA-centered structures have identified that such a method is not universally tractable for all listed species, is computationally data-intensive, and are constrained to the model’s time frame (*40,41*). Recovery plans are now structured on a conservation biology framework (“*The three R’s*”) focused on the ESA’s requirements of geographic representation of the species, conservation of the relevant ecosystems for the species to be self-sustaining, and abatement of threats (*42*). This conservation-based infrastructure should result in the establishment of multiple large, genetically-robust, self-sustaining populations across the species’ range and all ecological contexts.

As part of a recent effort for re-evaluating the red wolf recovery plan, the Association of Zoos and Aquariums’ American Red Wolf SAFE Program Action Plan (2019-2022) conducted a PVA and recommended that stakeholders work to ensure an *ex situ* population to support continued recovery efforts. Here, we defined significant levels of red wolf genomic ancestry in coyotes of southwestern Louisiana that are thriving on land where lethal management is not permitted. Over 50 years ago in this same geographic region is where the red wolf was last documented, subsequently declared extinct in the wild (*17,18,22,23,31,43-45*).

Given this high red wolf ancestry content along coastal SWLA, we suggest these coyote populations represent a potential for conservation redundancy of red wolf genes and the persistence of ancestral variation once thought to be extinct. As coastal SWLA is within the recent historic range of red wolves, including these populations in the red wolf’s three R’s recovery plan will promote the red wolves’ potential for adaptation, especially in a changing climate. Alongside PVA models, we suggest that conservation strategies include a mechanism to prioritize several aspects of admixed genomes (e.g. timing since admixture, percent content, ancestry block length) and thus the red wolf genomic legacies. These ghost genomes have naturally persisted in isolated areas for several decades through means not yet well understood.

We do acknowledge the challenge for implementation of a strict genomic ancestry profile. When possible, morphometrics are crucial for consideration of the ecological habitat in which the red wolf ancestry is found and the ultimate role of the canids in the ecosystem. This is especially important given landscape changes across the red wolves historic range. Our findings of higher red wolf ancestry proportions in SWLA may also be explained in part by the history of clear-cutting and livestock operations initiated in the late 1800s (*46*). Was early coyote-red wolf hybridization (*47*) a possible mechanism by which such canids were able to survive in a rapidly fragmented and converted landscape? Exclusion of individuals from conservation protection that do not conform to a phenotypic standard of an endangered species may result in the major oversight or exclusion of critical genomic variation potentially useful for genomic rescue or local adaptation through targeted practices.

As technology continues to provide innovative methods, the Gulf Coast canids also represent a critical biobanking opportunity for when genome editing methods are applied to red wolves. These methods were recently developed as a therapeutic technique to replace targeted gene sequences through the DNA repair process or transiently modify RNA (*48*). The consideration of such pioneering methods is the new frontier of conservation science for endangered species in the new era of anthropogenic-driven biodiversity loss and maladaptation due to rapidly changing climate and landscapes (*14,15*). We are at a pivotal moment where red wolves can be at the forefront to benefit from these developing conservation tools, and it is imperative to act quickly to preserve and harness red wolf ghost genomes now only present Gulf Coast canids.

## Materials and Methods

### Experimental Design

#### Sample Collection

From February–May 2021, we captured 26 coyotes using foothold traps with offset jaws (Minnesota Brand 550, Minnesota Trapline Products, Pennock, Minnesota, USA). Once captured, animals were restrained with a catchpole, muzzle, and hobbles. When needed, we chemically immobilized animals with an intramuscular injection of 1.3/kg ketamine HCl and 0.2 mg/kg xylazine HCl to inspect inside their mouths for injuries. We recorded sex, weight, and body measurements for all animals, and estimated age by tooth wear (*49,50*) (Table S2). We categorized animals ≥2 years as adults, 1–2 years old as juveniles, and less than 1 year old as pups. We collected 5mL of whole blood in Longmire buffer from the cephalic veins of captured coyotes and opportunistically sampled ear tissue from road-killed coyotes. All coyotes were fitted with Lotek LiteTrack Iridium 360 GPS collars (Lotek, Newmarket, Ontario, Canada). Our capture and handling of animals followed guidelines approved by the American Society of Mammalogists (2020) and were approved by the Institutional Animal Care and Use Committee at Michigan Technological University (#1677987-2). We plotted latitude and longitude for each sampled location using the *qmplot* function in the *ggmap* v3.0.0 R package (*51*).

#### DNA Extraction

We collected high molecular weight genomic DNA from whole blood or tissue from 36 coyotes sampled from Louisiana and 10 red wolves from North Carolina using the DNeasy Blood and Tissue Kit (Qiagen) and followed the manufacturer’s protocol for mammals. We quantified DNA concentration using the Qubit 2.0 fluorometer system and subsequently standardized DNA to 5ng/μL.

#### RAD sequencing and bioinformatic processing

We prepared 46 (2 samples were duplicated) genomic libraries for RADseq following a modified protocol (*52*). Briefly, we used the *Sbf1* restriction enzyme to digest genomic DNA and ligated a unique 8-bp barcoded biotinylated adapter to the resulting fragments. The barcode allows us to pool equal amounts of each DNA sample followed by random shearing to 400bp in a Covaris LE220. We used a Dynabeads M-280 streptavidin binding assay to enrich the pools for adapter ligated fragments, followed by a size selection for fragments 300-400bp in size and purification using Agencourt AMPure XP magnetic beads. The libraries were then prepared for Illumina NovaSeq 2×150nt sequencing at Princeton University’s Lewis Sigler Genomics Institute core facility using the NEBnext Ultra II DNA Library Prep Kit.

We retained sequencing reads that contained the unique barcode and the remnant *SbfI* cut site. We processed read data in *STACKS* v2 to first demultiplex the pools using 2bp mismatch for barcode rescue in the *process_radtags* module. We retained reads with a quality score ≥10 and removed PCR duplicates with the paired-end sequencing filtering option with the *clone_filter* module. Cleaned reads were then mapped to the dog genome CanFam3.1 assembly (*53*) using *BWA-mem* (*54*). We also filtered mapped reads for a minimum MAPQ of 20 and converted to bam format in *Samtools* v0.1.18 (*55*). We included RADseq data from 88 canids that were previously published (coyotes=39, gray wolves=19, eastern wolves=10, captive red wolves=10) (Table S1). The 88 publicly available canid samples that were included as processed reads and mapped to the same reference genome assembly following these methods.

We completed SNP discovery using all samples to obtain a catalogue of all polymorphic sites possible. We followed the recommended pipeline for the *gstacks* and *populations* modules in *STACKS* v2 after the data was mapped to a reference genome (*56,57*). However, we increased the minimum significance threshold in *gstacks* to require more stringent confidence needed to identify a polymorphic site using the marukilow model (flags --*vt-alpha* and --*gt-alpha*, *p*=0.01). We reported all SNPs discovered per locus (opted against using the *populations* flag -- *write_single_snp*) as ancestry inference is best with high density data. We then used *VCFtools* v0.1.17 (*58*) to exclude singleton and private doubleton alleles, remove loci with more than 90% missing data across all samples, and removed individuals with more than 20% missing data (we excluded four samples, Table S1). We filtered for a minimum of 3% minor allele frequency (MAF) in *PLINK* v1.90b3i (*59*). For initial screening of the samples, we constructed a *“statistically neutral and unlinked”* dataset of SNPs by excluding sites within 50-SNP windows that exceeded genotype correlations of 0.5 (with the *PLINK* argument --*indep-pairwise* 50 5 0.5) and deviated from HWE with the argument --*hwe* 0.001. The PCA was completed in the program *flashPCA* (*60*).

#### Inclusion of 1970s canids from Texas

We included publicly available BAM files from 10 canid samples in the 1970s from Texas, previously mapped to the same reference genome assembly (*21*). Following the methods and thresholds detailed above, we annotated SNPs across 47 reference canids (37 coyotes and 10 red wolves), 10 canids captured during the 1970s, and the 31 coyotes from SWLA. Samples were excluded from downstream analyses if they contained at least 20% missing data.

### Statistical analysis

#### Inference of canid ancestry

We inferred local ancestry of 36 coyotes from Louisiana with possible red wolf ancestry with respect to two reference populations: coyotes and red wolves (defined in Table S1). Following our past methods, briefly, we implemented a two-layer hidden Markov model in the program *ELAI* to infer local genomic ancestry proportions for the 59,788 SNP set (*61*). We used the following parameters: -*C* set to 2 and -*c* set to 10. As the precise nature of admixture is unknown, we analyzed four time points since admixture (-*mg*): 5, 10, 15, and 20 generations. We implemented *ELAI* three times serially for each *-mg* parameter value with 30 EM steps and averaged results over all 12 independent analyses. *ELAI* returns a per-SNP allele dosage score, which estimates the most likely ancestry proportion. We assigned chromosomal positions with allele dosage between 0.8 and 1.8 as heterozygous and those with allele dosage >1.8 as homozygous.

#### Estimating the timing of admixture

We counted the number of ancestry block identity switches per individual genome. Given the reduced representation focus on *Sbf1* cut sites and size selection step, the resulting blocks are inflated in size. As such, admixture timing estimates are likely skewed towards more recent timing of admixture events. Following from Johnson and colleagues (*62*), we estimated the number of generations since admixture for diploid genomes from the equation:

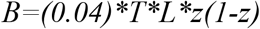

where *B* is the estimated number of ancestry switches, *T* is the number of generations since admixture, *L* is the total genome length (2085cM for autosomes and 111cM for the X chromosome; *63*), and *z* is the genome-wide red wolf ancestry proportion specific to autosomes or X chromosome. To convert the generation time into calendar years, we averaged the number of years since admixture across two generation times: the commonly estimated value of 4 years/generation and an estimate of 2 years/generation to account for scenarios in which a fraction of canids breed in their first year of life (*64,65*).

#### Morphology and red wolf ancestry

We assessed the relationship between ancestry estimates and body size with simple linear regression models with the lm function in program R. Response variable was body weight (kg), where we ran separate models with autosomal and X-linked red wolf ancestry estimates as explanatory variables and included sex in each model; all models fit a normal distribution. Models were evaluated for fit and adherence to assumptions by visualizing residuals and fitted values.

#### Maximum likelihood clustering method for population genetic structure analysis

We used the program *ADMIXTURE* (*66*) to assess proportional cluster membership (*Q*) across nine data partitions (*K*=2-10). We implemented the cross-validation (--*cv*) error flag to assess the best fit partition given the genotype data. Although the lowest *cv* error is presumed to be the best fit partition, we surveyed partitions with similar *cv* errors to evaluate the patterns of clustering with increasing partitions. Cluster patterns are likely influenced by relatedness and inbreeding, often an aspect of capture populations that is unavoidable (i.e. captive red wolf population).

## Acknowledgments

We are grateful for the contributions made by Bill Bruce and the FR Ranch of the Moore-Odom Wildlife Foundation, who provided us with access and support to study the coyotes on his property.

## Funding

This study was funded through supported received by KEB from the United States Fish and Wildlife Service, F20AC11140-00.

## Author contributions

Conceptualization: BMV, KEB, LPW

Methodology: BMV, JWH, KEB

Investigation: BMV, JWH, KEB

Visualization: BMV, AS

Writing—original draft: BMV, KEB, JWH

Writing—review & editing: BMV, KEB, JWH, LPW, JRA

## Competing interests

Authors declare that they have no competing interests.

## Data and materials availability

The 45 RADseq BAM files sequenced in this study have been submitted to the NCBI BioProject database (https://www.ncbi.nlm.nih.gov/bioproject/) under accession number PRJNA787606. See Table S1 for references to additional public data included in this study.

## Notes

### Competing Interest Statement

The authors have declared no competing interest.

